# Revisiting phage–host inference from metagenome-assembled genomes reveals the need for scaffold-level validation

**DOI:** 10.64898/2026.04.22.720271

**Authors:** Ling Yuan, Karthik Anantharaman, Bin Ma, Antonio Perdo Camargo, LinXing Chen

## Abstract

Accurate phage–host prediction underpins ecological interpretation and experimental validation of uncultivated phages, yet most current approaches rely heavily on metagenome-assembled genomes (MAGs). This reliance implicitly assumes that MAG-level quality metrics, such as completeness and contamination, ensure scaffold-level correctness. Here, we demonstrate that this assumption is frequently violated. Tetranucleotide frequency (TNF) analyses reveal that CRISPR-associated regions often diverge compositionally from their host genomes, increasing the risk of misbinning, while experimentally validated phage–host genome pairs show consistent TNF separation between phages and their hosts, challenging compositional binning assumptions. Through controlled simulations, we further show that the incorporation of substantial amounts of TNF-matched, non-host phage sequences into high-quality MAGs does not substantially alter standard completeness and contamination estimates, rendering such errors largely undetectable. Together, these results expose a systematic and underappreciated source of bias in MAG-based phage–host inference and highlight the need for explicit scaffold-level validation beyond conventional genome-quality metrics. We further propose a practical scaffold-level validation framework that integrates structural linkage and taxonomic consistency to improve the reliability of phage–host inference from MAGs.

## Introduction

Viruses are the most abundant biological entities on Earth and have been recognized as integral components of ecosystems across the planet ^1–3^. Viruses that exclusively infect bacteria are known as bacteriophages (or phages) ^4,5^. Phages exert profound influences on microbial communities by shaping community structure and dynamics through host infection and mortality ^6^, modulating biogeochemical cycling via auxiliary metabolic genes (AMGs) ^7,8^, and driving microbial evolution by horizontal gene transfer ^9,10^. Over the past 10–15 years, advances in metagenomics and viromics, together with the development of diverse analysis tools ^11–22^, have fundamentally reshaped our understanding of phages ^23^, revealing an immense and previously unrecognized diversity across a wide range of environments. Large-scale surveys have uncovered phages belonging to novel lineages ^24–28^, encoding diverse AMGs ^29–31^, as well as potential associations with human health ^32–34^. Together, these discoveries have broadened the prevailing view of phages, from being studied largely in pathogenic or host-centric contexts to being recognized as key ecological and evolutionary drivers in microbial ecosystems.

Accurate host prediction is not only central to ecological interpretation of phages but also plays a critical guiding role in their isolation, cultivation, and experimental characterization ^35^. The gold standard for host assignment remains isolation-based approaches, including plaque assays, infection experiments, and host range testing ^36^. However, the vast majority of phages discovered to date have been identified through metagenomics, leaving their hosts experimentally unknown. This disconnect between large-scale sequence discovery and laboratory validation has driven the widespread development and application of *in silico* host prediction methods, yet no single one of them is universally reliable ^37–39^. One reason for this limitation is that different methods rely on distinct principles and achieve varying levels of resolution. In addition, the reliability of the underlying reference databases represents a potentially significant but often overlooked source of uncertainty ^40,41^, especially the metagenome-assembled genomes (MAGs). MAGs form the basis of several widely used homology-based approaches corresponding to AMGs, tRNAs, and host-encoded CRISPR-Cas systems, as well as direct prediction of proviruses from MAGs. They have become increasingly available through public repositories ^40,42–45^ and occupy a central position among these strategies because they serve as the common reference framework for multiple host prediction signals by providing genome-resolved representations of uncultivated bacteria ^41^. This widespread reliance on MAGs implicitly assumes that MAG-level genome quality metrics (primarily completeness and contamination) are sufficient to ensure the correctness of host prediction at the scaffold level. However, because mobile genetic element contigs are disproportionately misbinned, MAG-based host inference may be systematically biased, and the magnitude of this problem has not been evaluated in a standardized way ^46^.

Here, we systematically evaluate the reliability of MAG-based phage–host relationship prediction by examining the key assumptions that underpin current genome-resolved metagenomic workflows. Specifically, we test whether scaffold-level features commonly used for host inference, including CRISPR–Cas loci and viral sequences, exhibit compositional profiles consistent with those of their assigned MAGs. We further assess whether standard MAG quality metrics are sufficiently sensitive to detect the incorporation of unrelated viral scaffolds. By combining TNF-based compositional analyses of CRISPR-associated regions, experimentally validated host–phage genome pairs, and controlled simulations of viral scaffold incorporation into high-quality MAGs, we demonstrate that scaffold-level misassignment can remain largely invisible to conventional genome-quality assessments. Our results reveal an underappreciated and potentially widespread source of bias in MAG-based phage–host prediction and highlight the need for explicit scaffold-level validation in both newly reconstructed and publicly available MAG datasets.

## Results

### Issues in using metagenome-assembled genomes for phage-host relationship prediction

Metagenome-assembled genomes (MAGs) consist of one or multiple contigs or scaffolds that are grouped based on their sequence composition (usually assessed through tetranucleotide frequencies, TNF) and sequencing coverage profiles (Figure 1A) ^47^. The quality of MAGs is commonly assessed using metrics such as completeness and contamination, together with auxiliary features including the presence of ribosomal RNA (rRNA; 23S, 16S, and 5S) and tRNA genes ^48^. Importantly, these metrics are primarily derived from conserved host genomic features, such as single-copy marker genes, and do not explicitly incorporate phage-related signals. As a result, they are inherently unable to evaluate whether viral scaffolds have been incorrectly incorporated into a MAG. MAGs passing stringent thresholds of completeness (for example, >90%) and contamination (for example, <5%) are often treated as reliable representations in downstream analyses, including the prediction of phage-host relationships. However, these MAG-level quality metrics do not necessarily guarantee scaffold-level correctness. Our previous analyses demonstrated that MAGs with low estimated contamination can still contain unrelated scaffolds originating from distinct organisms because of coincidentally similar TNF and/or sequencing coverage ^49^. As a result, the presence of a scaffold encoding a CRISPR–Cas repeat region derived from another organism, or the inclusion of a viral scaffold whose true host differs from that of the MAG, may have little to no effect on the overall completeness and/or contamination estimates of the MAG. We argue that the inclusion of unrelated CRISPR-Cas and/or viral scaffolds will strongly affect the accuracy of downstream phage-host relationship prediction (Figures 1B and C). In principle, some of these misbinning events can be mitigated by incorporating sequencing coverage profiles from multiple related samples during genome binning. However, this information is not always available or sufficient. Alternatively, unrelated scaffolds can be identified and removed through post-binning genome curation, a step that is often omitted in routine metagenomic workflows but is critical for ensuring the reliability of phage–host relationship inference (Figure 1A).

**Figure 1.**
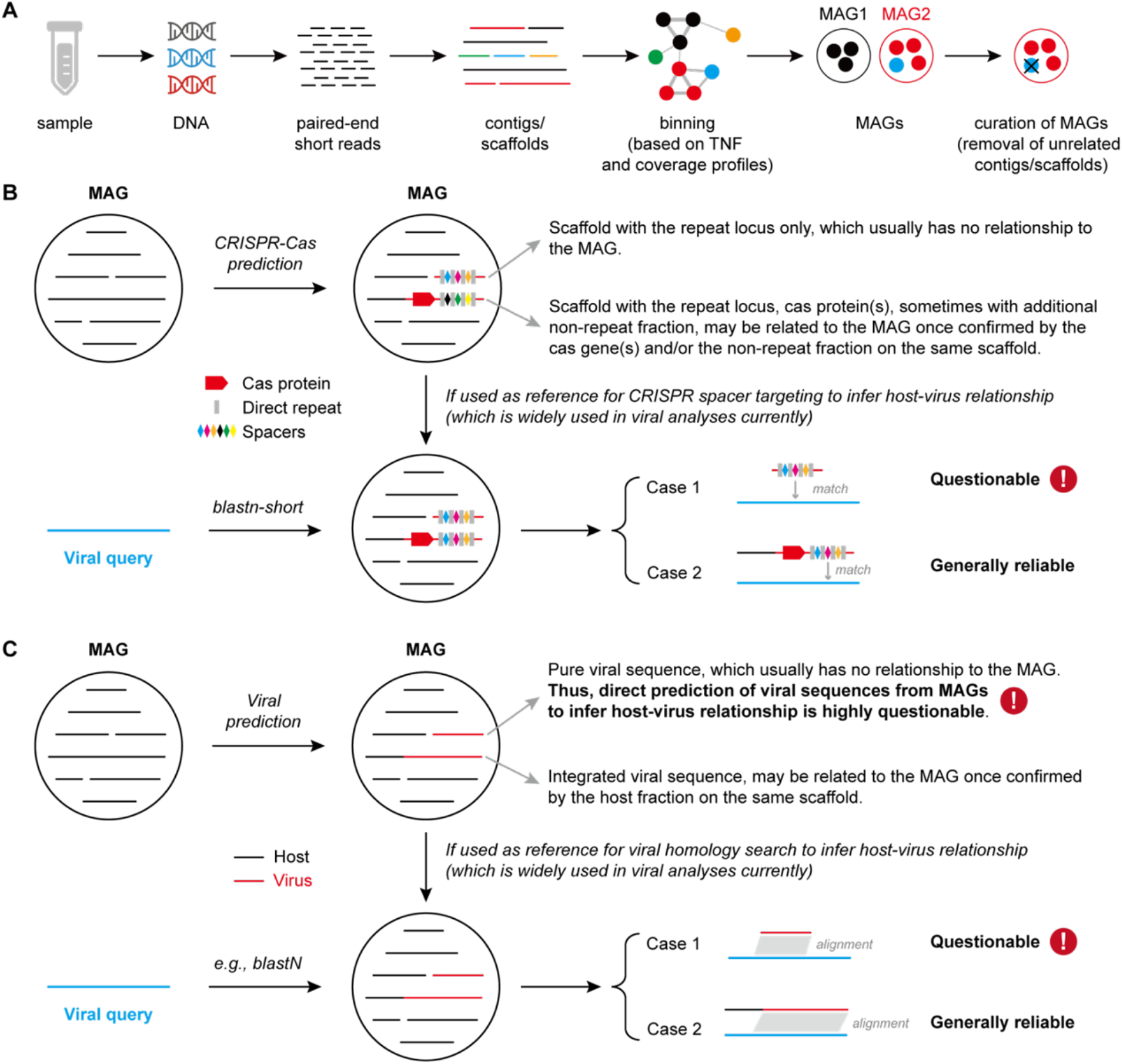
Concept illustration of the common pitfalls in using MAGs to infer phage-host relationships. (A) General workflow of metagenomic analyses leading to the reconstruction of metagenome-assembled genomes (MAGs). The curation of MAGs represents a critical step for removing unrelated contigs or scaffolds. However, this step is frequently omitted in routine analyses. (B) A MAG may contain a scaffold with only the CRISPR repeat locus region (repeats + spacers), which is usually from another unrelated MAG. For a scaffold with a CRISPR repeat locus, cas gene(s), and additional host fraction, the longer the host fraction is, the more reliable the confirmation can be. (C) Viral scaffolds in MAGs may have no relationship to each other. For an integrated viral sequence, the longer the host fraction is, the more reliable the confirmation could be. In the metagenomic analyses based on short-reads sequencing and assembly, the assembled scaffolds are generally binned to obtain MAGs based on their TNF and/or sequencing coverage profiles. However, the scaffolds from infecting viruses and the CRISPR repeat locus regions usually do not have consistent TNF and/or coverage with those of the hosts.

To systematically examine potential sources of error in phage-host prediction based on MAGs, we summarized the most common analytical scenarios in which MAGs are used to infer phage-host associations (Figures 1B and C). These analyses include (1) linking viruses to MAGs through matches to CRISPR–Cas spacers detected on MAG contigs (Figure 1B), (2) identifying viral scaffolds directly within MAGs and assigning the MAG taxon as the viral host, or inferring host identity based on sequence homology between viral scaffolds and MAG scaffolds (Figure 1C). In practice, these analyses frequently rely on MAGs obtained either from public genome repositories (e.g., NCBI, GTDB) or from the same metagenomic datasets used for viral identification. These scenarios lead to multiple modes of erroneous phage-host association, including the assignment of a virus to an incorrect host, or the inference of host identity based on misassigned CRISPR–Cas repeat loci. Importantly, such errors cannot be detected by standard MAG quality metrics alone, highlighting a fundamental limitation of MAG-based phage-host relationship prediction when scaffold-level validation is not explicitly performed.

### CRISPR-encoding regions exhibit distinct sequence composition from their host genomes and may fail to co-bin into correct MAGs

CRISPR–Cas systems constitute adaptive immune mechanisms in bacteria and archaea that provide defense against invasive genetic elements, including viruses and plasmids ^50^. Within a given microbial population, strains sharing the same CRISPR–Cas system can nevertheless differ substantially in their repeat–spacer loci, primarily due to the continuous acquisition of new spacers ^51,52^. In metagenomic assemblies based on short paired-end reads, these highly variable CRISPR–Cas loci are therefore frequently fragmented, resulting in multiple scaffolds that encode different components of the system (Figures 2A and 2B). In addition, CRISPR– Cas systems are known to undergo frequent horizontal gene transfer ^53^, raising the possibility that CRISPR-encoding regions may differ in sequence composition from the rest of the host genome and thus fail to co-bin into the same MAG.

**Figure 2.**
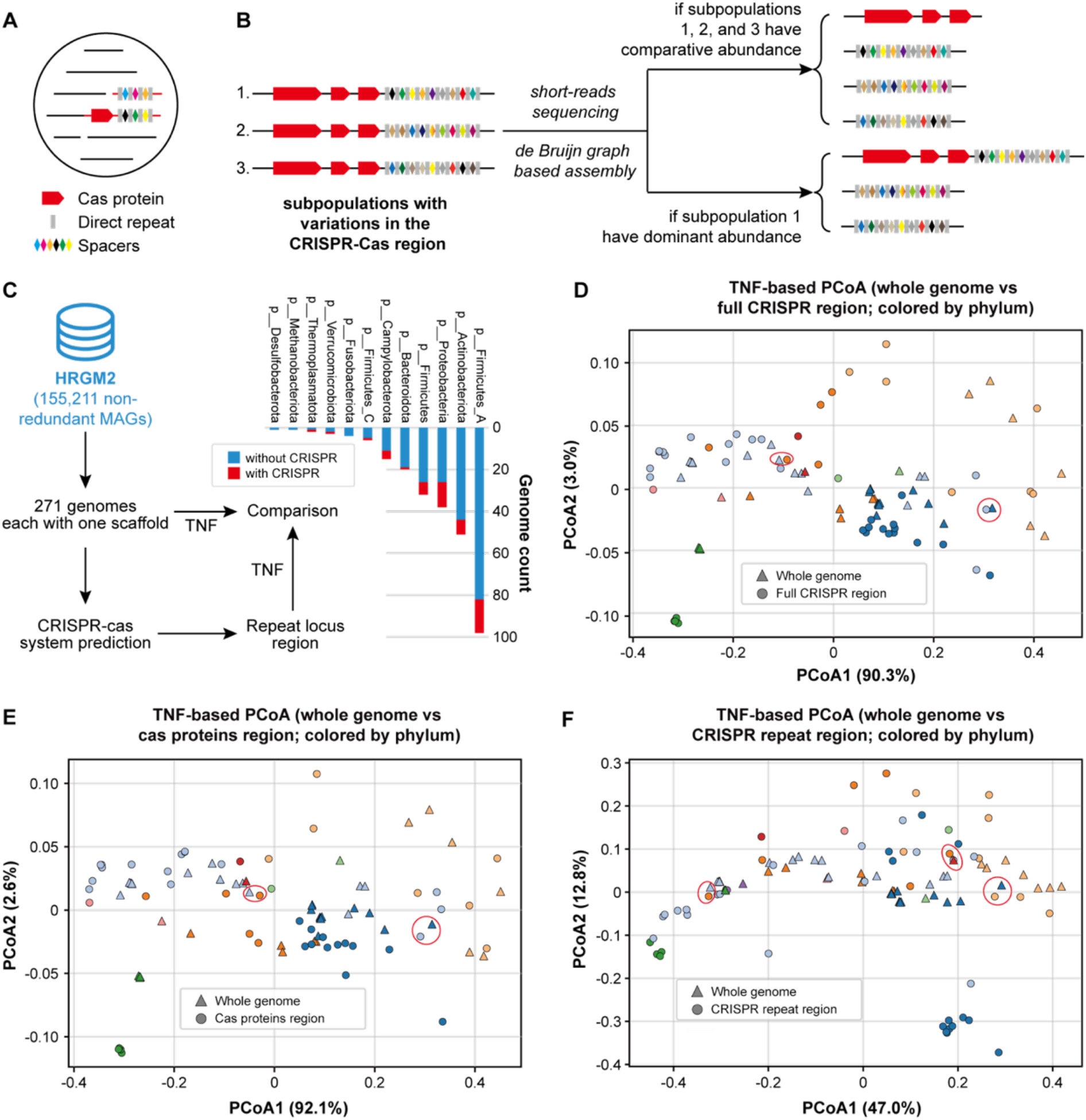
CRISPR–Cas–encoding scaffolds in metagenomic analyses. (A) Schematic illustration of the two principal modes by which CRISPR–Cas–encoding scaffolds are incorporated into metagenome-assembled genomes (MAGs). (B) Representative scenarios illustrating how genomic fragments are generated during metagenomic assembly of short paired-end reads using de Bruijn graph–based assemblers. (C) Identification and characterization of CRISPR–Cas systems within 271 single-scaffold MAGs from the HRGM2 dataset (Supplementary Table 1). TNF–based PCoA (based on a cosine distance matrix) comparing sequences of the whole genomes against (D) the full CRISPR-Cas system regions, (E) the Cas protein-coding regions, and (F) the CRISPR repeat regions. Red circles indicate sequences of whole genomes and CRISPR-related regions from different phyla, which could be binned together if they were with comparative sequencing depth and presented in the same community.

To evaluate this possibility, we identified 271 single-scaffold MAGs from the HRGM2 database ^45^ and detected a total of 48 CRISPR–Cas systems encoded within these genomes (Figure 2C). We then compared the sequence composition of whole genomes with that of CRISPR-associated regions using TNF–based principal coordinates analysis (PCoA). Specifically, we performed separate ordination analyses comparing whole genomes against the full CRISPR–Cas regions (Figure 2D), the regions encoding Cas proteins (Figure 2E), and the CRISPR repeat array regions alone (Figure 2F). Although both whole genomes and CRISPR-associated regions were colored according to their phylum-level taxonomic assignments, CRISPR-related sequences, particularly the CRISPR repeat regions, often occupied distinct regions of TNF space relative to their corresponding host genomes. In several cases, CRISPR-derived sequences were compositionally closer to sequences from different phyla than to their own host genomes. The same phenomenon was observed at the genome level for those from Firmicutes_A and Pseudomonadota (Supplementary Figure 1). These results indicate that CRISPR-associated scaffolds can exhibit pronounced compositional divergence from the rest of the genome.

These observations suggest that CRISPR-encoding scaffolds are at high risk of being mis-binned in metagenomic analyses, producing MAGs that contain contigs with CRISPR spacers originating from genomes other than the one represented by the MAG. Such mis-binning could in turn lead to erroneous inference of host–virus relationships if MAGs are used as the source for CRISPR spacers used for establishing putative phage-host relationships.

**Figure S1. (Figure 2 related).**
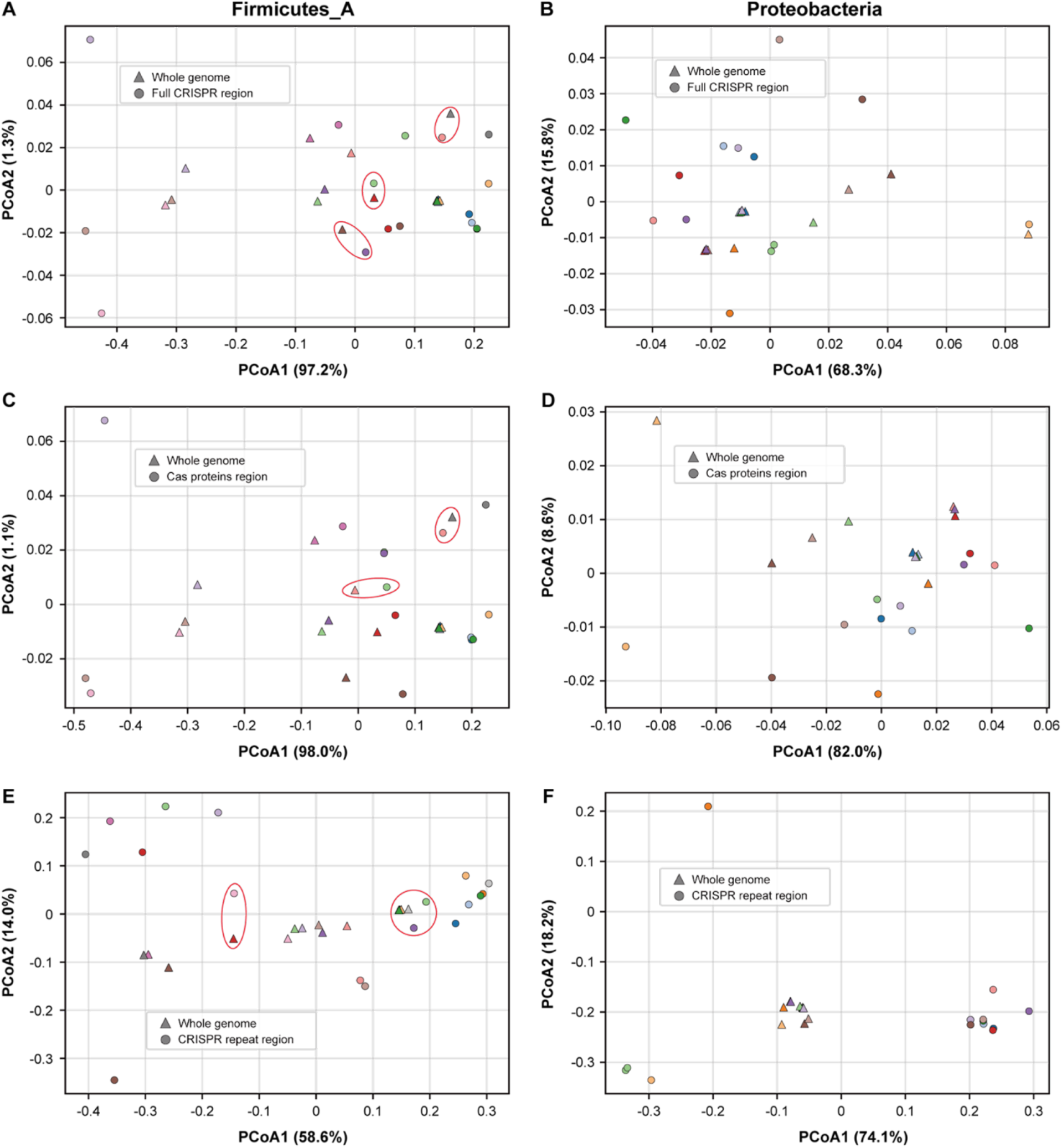
TNF–based PCoA comparison of CRISPR-related sequences against the whole genomes at the genome level. Comparison of the whole genomes against (A) and (B) the full CRISPR-cas system regions, (C) and (D) the cas proteins regions, (E) and (F) the CRISPR repeat regions, for those from the phylum of (A) (C) (E) Firmicutes_A and (B) (D) (F) Pseudomonadota. Red circles indicate sequences of whole genomes and CRISPR-related regions from different genome, which could be binned together if with comparative sequencing depth and presented in the same community.

### Viruses may exhibit distinct sequence composition from their hosts and are therefore unlikely to be co-binned in metagenomic analyses

To assess whether bacterial hosts and their phages exhibit systematic differences in sequence composition, we calculated genome-level TNF for host genomes and their experimentally validated infecting phages and visualized their relationships using PCoA. Genomes from *Streptomyces* (phylum Actinobacteriota) and *Escherichia coli* (phylum Pseudomonadota), together with their corresponding phage genomes, were used as representative examples (Figures 3A and B, Supplementary Tables 2 and 3).

**Figure 3.**
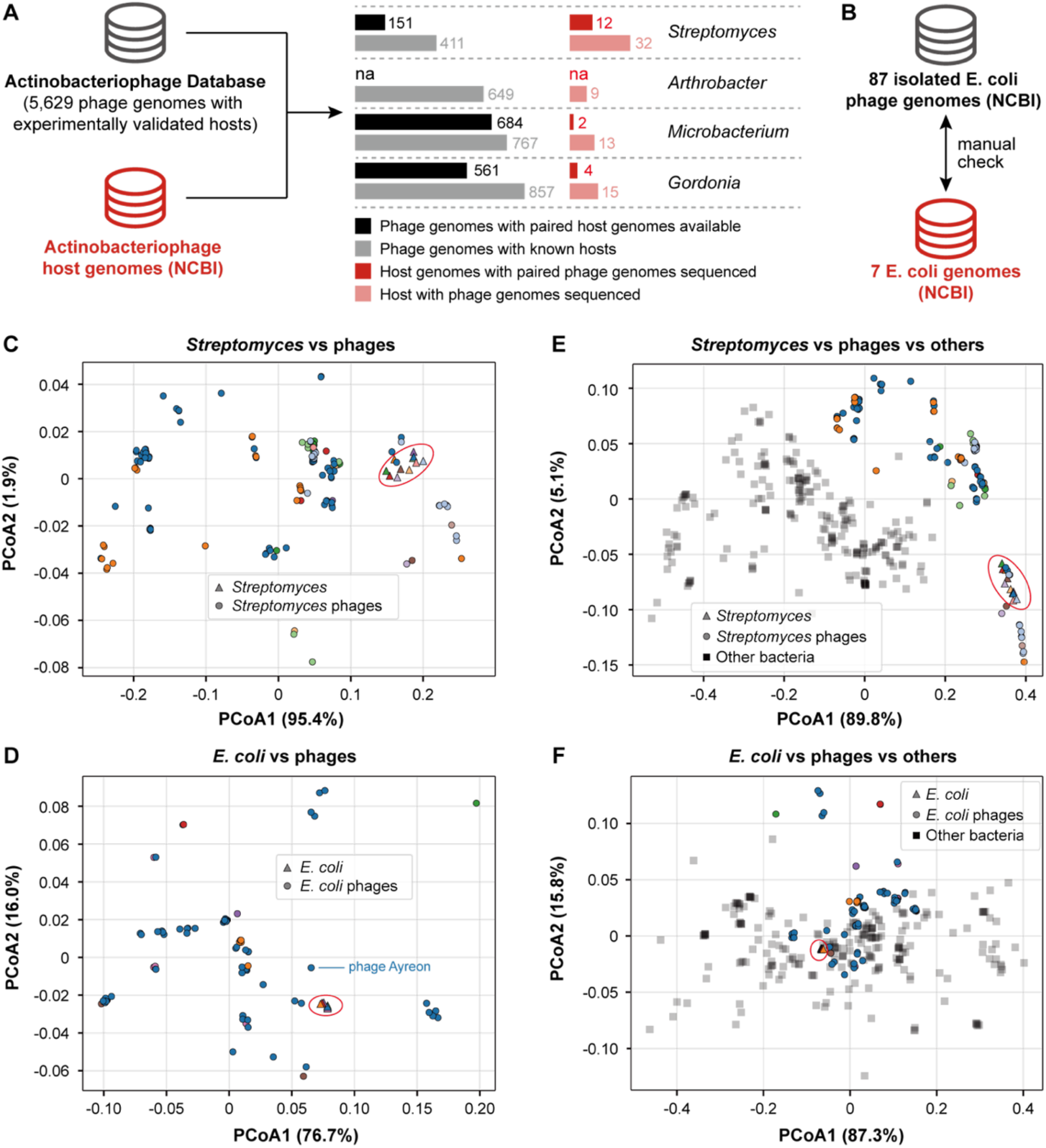
Phage genomes exhibit TNF compositions distinct from their bacterial hosts. (A) Overview of experimentally validated actinobacteriophage–host associations curated from the Actinobacteriophage Database ^54^. The four host genera with the largest numbers of sequenced infecting phage genomes are shown (Supplementary Table 2). (B) Manual retrieval of genomes for *E*.*coli* and their isolated phages from NCBI (Supplementary Table 3). Principal coordinates analysis (PCoA) of genome-level TNF profiles for host genomes (scaffolds ≥ 5 kbp) and their experimentally validated infecting phage genomes of (C) *Streptomyces* spp. and (D) *E. coli*. TNF-based PCoA analysis of (E) *Streptomyces* and their phage genomes with 220 non-Actinobacteriota single-scaffold bacterial MAGs (Supplementary Table 1), and (F) *E. coli* and their phage phages along with 233 non-Pseudomonadota single-scaffold bacterial MAGs (Supplementary Table 1). In (C)-(F), host genomes were indicated by triangles, phage genomes by circles, and other bacterial MAGs by squares. Host–phage pairs are indicated using identical colors. The host genomes are highlighted by the red circles.

In both cases, host and phage genomes occupied largely distinct regions of TNF space (Figures 3C and D). Host genomes formed compact and well-defined clusters, whereas phage genomes were generally displaced away from their corresponding hosts. When additional bacterial MAGs from outside the host phylum were included, several phage genomes exhibited greater compositional similarity to non-host bacteria than to their confirmed hosts (Figures 3E and F). These patterns reveal two notable features of the TNF landscape. First, some similar phages infecting the same host tend to occupy related regions of compositional space, reflecting shared constraints or evolutionary history. Second, despite this association, most phage genomes remain compositionally distinct from their host genomes at the tetranucleotide level and can, in some cases, more closely resemble unrelated bacterial genomes.

Collectively, these results demonstrate a pronounced and consistent divergence in tetranucleotide composition between bacterial hosts and their phages, even among experimentally confirmed host–phage pairs. This finding highlights a fundamental limitation of TNF-based similarity as a criterion for linking phages to their hosts in metagenomic binning. Moreover, this challenge may be further exacerbated by the fact that the sequencing coverage of virulent phages does not necessarily mirror that of their hosts, particularly during active lytic replication or in the presence of abundant extracellular viral particles.

Together, these observations underscore the need for caution when relying solely on compositional and coverage-based signals for host-phage association inference.

### Limited sensitivity of MAG quality metrics to the incorporation of compositionally similar but non-host viral sequences

To test whether incorporation of viral sequences can substantially affect commonly used MAG quality estimates, we performed a controlled simulation in which Actinobacteriophage-derived scaffolds were incorporated into high-quality bacterial (non-Actinobacteriota) MAGs based on TNF similarity (Figure 4A; see Methods). The resulting modified MAGs were then re-assessed using CheckM and CheckM2, and completeness and contamination estimates were compared with those of the original MAGs (Figures 4B-E).

**Figure 4.**
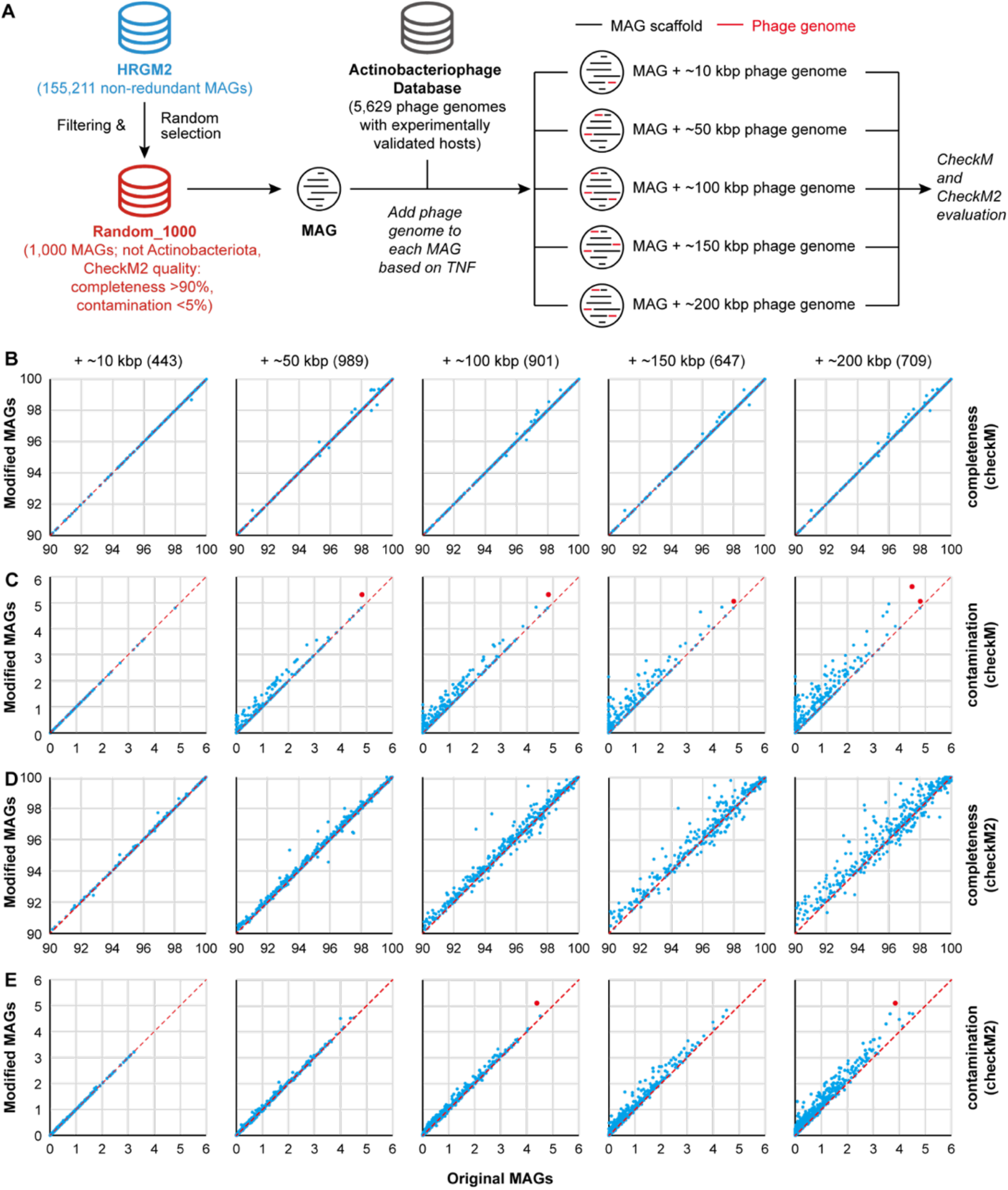
Incorporation of compositionally similar but non-host viral scaffolds into MAGs minimally affects completeness and contamination estimates. (A) Selection of high-quality bacterial MAGs and Actinobacteriophage genomes used for simulation. A total of 1,000 high-quality bacterial MAGs (completeness ≥ 90% and contamination ≤ 5% by CheckM2), excluding Actinobacteriota, were randomly selected from the HRGM2 genome collection ^45^. Actinobacteriophage genomes were obtained from the Actinobacteriophage Database ^54^ and used as viral sequence sources. For each MAG, viral genomic sequences with predefined total lengths (approximately 10 kbp, 50 kbp, 100 kbp, 150 kbp, or 200 kbp) were incorporated based on TNF similarity. Some MAGs were excluded from downstream analyses because no suitable viral sequences with sufficiently similar TNF profiles could be identified (see Methods). (B)-(E) Comparisons of completeness (B and D) and contamination (C and E) between original and modified MAGs using CheckM (B and C) and CheckM2 (D and E). In panels (B)-(E), each point represents one MAG, and the numbers of original–modified MAG pairs are indicated in brackets. The dashed red line indicates y = x. In panels (C) and (E), the five and two modified MAGs with contamination estimates exceeding 5% are highlighted with red circles.

As evaluated by CheckM, completeness showed minimal deviation following viral sequence incorporation across all simulated conditions, with a strong correspondence between original and modified MAGs along the identity line (y = x), even when up to ∼200 kbp of viral sequence was added (Figure 4B). In contrast, CheckM2 revealed a bidirectional shift in completeness, and the magnitude of deviation increased with the total length of incorporated viral sequences (Figure 4D).

Contamination increased and scaled positively with the amount of viral sequence added according to both CheckM and CheckM2 (Figures 4C and E). Nevertheless, contamination estimates for the vast majority of MAGs remained below the commonly used 5% threshold, with only five (CheckM) and two (CheckM2) modified MAGs exceeding 5% contamination under the simulated conditions.

Together, these results indicate that standard completeness and contamination metrics exhibit only limited sensitivity to the incorporation of substantial amounts of TNF-matched, non-host viral sequences. Detectable increases in contamination were largely restricted to higher incorporation levels (the ≥ 50 kbp simulation by CheckM, Figure 4C; and the ≥ 100 kbp simulation by CheckM2, Figure 4E), whereas smaller amounts of viral sequence had little effect on the contamination scores. This limited sensitivity implies that high completeness and low contamination alone are not sufficient to rule out the presence of non-host viral sequences in MAGs, particularly in workflows that rely heavily on compositional similarity for binning and quality assessment. Notably, identifying and removing viral contaminants from MAGs would be expected to reduce contamination both in the underlying assemblies and in marker-based quality estimates.

### A scaffold-level validation framework for improving phage–host inference from MAGs

Based on the systematic biases revealed above, we next sought to define a set of practical criteria for improving the reliability of MAG-based phage–host inference by explicitly incorporating scaffold-level validation. We acknowledge that these are practical for archaeal virus analyses as well. Because viral and CRISPR-associated sequences frequently violate the compositional and coverage assumptions underlying genome binning (Figures 2–4), we reasoned that host assignment should not rely solely on the co-occurrence of viral or CRISPR signals within a MAG, but instead require explicit evidence linking these signals to the host genome (i.e., MAG) at the scaffold level (Figure 5).

Firstly, we should distinguish between integrated and non-integrated viral scaffolds identified within MAGs. Viral scaffolds lacking evidence of integration, e.g., absence of host–virus junctions or contiguous host genomic regions, are not considered sufficient to support phage–host association, as such sequences may represent co-binned but unrelated viral genomes.

Second, for candidate integrated viral scaffolds, we should require the presence of a sufficient length of flanking host-derived sequence to establish genomic context. Specifically, scaffolds are retained only if they encode multiple non-viral genes (at least five, preferably 10 or more), whose taxonomic assignments, inferred using a lowest common ancestor (LCA) approach, should be consistent with the taxonomy of the corresponding MAG as determined by GTDB-Tk or other comparable tools, at a minimum resolution of the family level. This criterion ensures that the viral sequence is reliably linked to a host genomic background rather than co-binned based on coincidental compositional similarity, such as TNF.

Thirdly, similar considerations should be applied to scaffolds containing CRISPR–Cas systems identified within MAGs. Scaffolds containing only CRISPR repeat–spacer arrays, without associated cas genes or additional host genomic context, should be excluded from downstream analyses, as such sequences are particularly prone to fragmentation and misbinning.

Fourth, for CRISPR-associated scaffolds that include cas genes and additional non-cas regions, we again require the presence of at least five, preferably 10 or more such genes, whose taxonomic assignments are concordant with the MAG at the family level or lower, as described above. This filtering step aims to ensure that CRISPR spacers used for host–virus linkage originate from the corresponding host of the MAG, rather than from any unrelated taxa.

In summary, these criteria define a scaffold-level validation framework that integrates structural linkage and taxonomic consistency to reduce false-positive virus–host associations arising from MAG misbinning.

Application of such constraints is expected to substantially improve the robustness of host inference in both newly reconstructed and publicly available MAG datasets, particularly in analyses that rely on the identification of viral scaffolds within MAGs or CRISPR spacer matching.

## Discussion

As the majority of the phages could not be studied in cultivation, bioinformatic approaches are widely used to infer their bacterial hosts, which are heavily based on MAGs (^37–39^). Here, we demonstrate that MAG-level quality metrics (such as completeness and contamination) ensure scaffold-level correctness, a central assumption underlying MAG-based phage–host relationship prediction, is frequently unjustified (Figures 1-4). Although completeness and contamination estimates have become standard criteria for evaluating MAG quality, these metrics were developed primarily to assess the reconstruction of cellular genomes such as bacteria and archaea ^55,56^, and are intrinsically insensitive to the incorporation of viral scaffolds or horizontally transferred mobile genetic elements ^57^. As a result, MAGs that satisfy stringent quality thresholds may nevertheless contain scaffolds that originate from unrelated organisms or viruses, leading to systematic errors in downstream host assignment.

A key finding of this study is that both CRISPR-associated regions (Figure 2 and Supplementary Figure S1) and experimentally validated phage genomes (Figure 3) often exhibit TNF profiles that are markedly distinct from those of their corresponding hosts. This observation challenges a fundamental premise of composition-based genome binning, in which biologically linked sequences are expected to occupy similar compositional space. For CRISPR–Cas loci, this divergence is likely driven by a combination of rapid spacer turnover, local repeat enrichment, and potential horizontal transfer events ^58–60^. For phages, the divergence reflects their independent evolutionary histories, genome architectures, and selective constraints. Importantly, in some cases, phage genomes were compositionally closer to unrelated bacterial genomes than to their experimentally confirmed hosts (Figure 3), highlighting the risk that TNF-based binning may incorrectly associate viral scaffolds with non-host MAGs. Coverage-based binning strategies could further compound this problem. Unlike host chromosomes, virulent phages do not necessarily exhibit sequencing coverage profiles mirroring those of their bacterial hosts. During active lytic replication, intracellular amplification and the accumulation of extracellular viral particles can substantially inflate viral coverage relative to the abundance of the corresponding hosts. Conversely, prophages in low-copy or partially induced states may display discordant coverage patterns across samples. These dynamics reduce the reliability of co-abundance as an independent signal for host assignment. In summary, due to these natural features of phages, the two principal dimensions used in conventional binning, i.e., sequence composition and sequencing coverage, may both be systematically violated for viral sequences.

Our simulation analyses further demonstrate that even substantial incorporation of TNF-matched, non-host viral sequences can have minimal effects on standard completeness and contamination estimates (Figure 4). This finding is particularly important because it shows that MAG-level quality metrics are not merely imperfect but fundamentally poorly aligned with the problem of viral misbinning. In practice, this means that high-quality MAGs deposited in public repositories may still contain misassigned viral scaffolds that remain undetected under current quality-control standards. Such hidden errors may propagate through downstream analyses that rely on reference MAGs, including CRISPR spacer matching, prophage identification, AMG interpretation, and host prediction frameworks based on machine learning. For example, a recent study reported the identification of thousands of lytic phages in bacterial genome assemblies, which challenged current assumptions regarding the nature of the lytic lifestyle of phages ^61^. Because the analyzed dataset was derived from large-scale RefSeq bacterial genome assemblies, which likely include metagenome-derived genomes and other non-isolate reconstructions, it remains unclear to what extent some of the identified lytic phage signals may reflect scaffold-level misbinning or assembly artifacts.

The implications of these findings extend beyond phage–host prediction. Mobile genetic elements (MGEs), including plasmids, genomic islands, integrative conjugative elements, and transposon-rich regions, are all prone to scaffold-level misbinning in short-read metagenomic assemblies ^49^. Therefore, the issue identified here likely reflects a broader methodological limitation in genome-resolved metagenomics. As MAG repositories continue to expand and increasingly serve as foundational reference resources for ecological and evolutionary inference, the scaffold-level integrity of these genomes warrants much greater attention when studying MGEs.

Taken together, our findings argue for a shift from MAG-centric confidence to scaffold-aware validation in phage–host relationship inference workflows. We suggest that future analyses should incorporate explicit post-binning curation to confirm the validation of CRISPR-Cas and virus-related sequences in both public and self-reconstructed MAGs, or apply orthogonal host prediction signals such as experimentally supported CRISPR spacer matches. By highlighting these overlooked pitfalls, this work provides a methodological framework for improving the robustness of phage–host relationship prediction in the metagenomic era.

## Methods

### TNF-based comparison of CRISPR repeat regions and the corresponding whole bacterial genomes

To compare sequence composition between whole bacterial genomes and CRISPR repeat regions, we performed a TNF–based ordination analysis. We identified a total of 271 single-scaffold MAGs from the HRGM2 datasets ^45^, and the CRISPR-Cas systems were predicted from them. Specifically, we first predicted the CRISPR repeat regions using PILER-CR version 1.06 with default parameters ^62^. Then, the protein-coding genes within 2,000 bp upstream and downstream of the predicted repeat regions were searched for cas proteins using hmmsearch against the TIGRFAM HMM database with suggested threshold scores. Only those repeat regions with at least one cas protein identified were counted as true CRISPR-Cas systems.

For each sequence of the bacterial genomes and the predicted CRISPR repeat regions, canonical tetranucleotide frequencies were calculated using a sliding window approach. All possible 4-mers composed of A, C, G, and T were considered, and each k-mer was collapsed with its reverse complement into a single canonical representation. Tetranucleotide counts were normalized by the total number of valid 4-mers in each sequence to obtain relative frequency profiles. No minimum length filtering was applied to CRISPR repeat array regions to retain short but compositionally informative sequences.

Pairwise compositional dissimilarities between sequences were calculated using cosine distance on the normalized TNF vectors. PCoA analysis was then performed on the resulting distance matrix using classical multidimensional scaling to project sequences into a low-dimensional compositional space. The proportion of variance explained by each axis was calculated from the eigenvalues of the centered distance matrix.

### TNF-based comparison of experimentally validated actinobacteriophage–host genome pairs

To evaluate whether viral genomes and their hosts exhibit systematic differences in TNF composition, we focused on actinobacteriophages that have experimentally validated host assignments. Host–phage relationships were obtained from the Actinobacteriophage Database ^54^, in which phage–host pairs are supported by isolation and infection experiments, thereby avoiding uncertainties associated with computational host prediction using bioinformatic tools.

We first identified the four host genera within the Actinobacteriophage Database (https://phagesdb.org/) that had the largest numbers of sequenced actinobacteriophage genomes. The corresponding host genomes were retrieved from NCBI when available. Among these four genera, only *Streptomyces* was represented by a sufficiently large number of publicly available host genomes with paired phage genomes to enable genome-level compositional analysis. Specifically, a total of 12 *Streptomyces* host genomes and 151 corresponding *Streptomyces*-infecting phage genomes were included (Supplementary Table 2).

We also searched NCBI using the keyword “E. coli phage” for isolated phage genomes of *E. coli*, and the corresponding host genomes were manually confirmed according to the NCBI descriptions of each phage genome. The host and isolated phage genomes were subsequently downloaded from NCBI when available, resulting in 7 *E. coli* and 87 phage genomes (Supplementary Table 3).

In addition, we added single-scaffold bacterial MAGs, the same ones we used in the CRISPR TNF-based analyses (Supplementary Table 1), to the *Streptomyces* and phage genome set, or the *E. coli* and phage genomes set, after removing the ones belonging to the same phylum.

For each host genome, scaffolds with a minimum length of 5 kbp were retained to minimize compositional noise associated with short sequences. TNF were then calculated at the genome level by aggregating all retained scaffolds from each host genome. For phages, TNF was calculated individually for each phage genome, with each genome treated as an individual analytical unit. TNF were computed as relative frequencies of canonical tetranucleotides, in which each 4-mer was merged with its reverse complement, and only unambiguous nucleotides (i.e., A, C, G, and T) were considered. Pairwise compositional distances between host and phage TNF profiles were calculated using cosine distance, and the resulting distance matrix was subjected to principal coordinates analysis (PCoA). The host and their phage genomes were projected into a shared ordination space to visualize compositional relationships.

### Evaluation of the incorporation of compositionally similar but non-host viral sequences into MAGs

To assess the impact of co-binned and unrelated viral sequences on the completeness and contamination of MAGs, we performed two simulations and evaluations, including (1) co-binning random viral sequences into 1000 MAGs, and (2) co-binning random viral sequences into the E. coli K12 MG1655 genome.

We first simulated the co-binning of random viral sequences into multiple MAGs. Briefly, 1,000 bacterial MAGs, not from Actinobacteriota and with >90% completeness and <5% contamination (evaluated by checkM2), were randomly selected from the 155,211non-redundant genome collection of HRGM2 ^45^. The information on the 1000 selected MAGs is available in Supplementary Table 4. The viral sequences were from the Actinobacteriophage Database (https://phagesdb.org/data/), which contains 5,629 genomes (downloaded on December 12, 2025) with experimentally validated hosts ^54^.

TNF of each Actinobacteriophage genome was calculated, and the average TNF of all scaffolds in each MAG was computed to represent the compositional profile of the corresponding MAG. Canonical TNF was used, in which each 4-mer was merged with its reverse complement, resulting in a 136-dimensional feature vector for each genome. For each MAG, the cosine similarity was calculated between its TNF profile and those of all Actinobacteriophage genomes. Based on these similarities, the top 100 most similar Actinobacteriophage genomes were selected as candidate viruses for that MAG. The Actinobacteriophage sequences were then stochastically introduced into each MAG by sampling from the candidate pool using a similarity-weighted scheme, in which the sampling probability of each Actinobacteriophage was proportional to a softmax transformation of its cosine similarity score. This approach preferentially selects viruses with TNF profiles more similar to the target MAG while retaining stochasticity in the selection process. For each MAG, one or more Actinobacteriophage genomes were added such that the total length of the introduced viral sequences fell within the define range, i.e., 10 kbp, 50 kbp, 100 kbp, and 200 kbp, within a tolerance of ±10 kbp. Each viral Actinobacteriophage was added at most once to a given MAG but could be reused across different MAGs. This procedure generated a set of simulated MAGs with controlled levels of viral sequence incorporation while preserving compositional similarity between host and viral sequences. For each modified MAG, metadata including original genome length and sequence count, number and name of added viral contigs, total added viral length, and final genome size were recorded in Supplementary Table 4.

For all the original MAGs and genomes and the simulated genomes, we evaluated their quality using CheckM version 1.2.4 ^55^ with the “lineage_wf” mode, and CheckM2 version 0.1.3 ^56^ with the “predict” mode, and the values of completeness and contamination were compared accordingly.

## Notes

### Competing Interest Statement

The authors have declared no competing interest.

